# The investigation of dysregulated visual perceptual organization in adults with autism spectrum disorders with phase-amplitude coupling and directed connectivity

**DOI:** 10.1101/2024.01.25.577004

**Authors:** Limin Sun

## Abstract

Phase-amplitude coupling (PAC) has been used as a powerful tool to understand the mechanism underlying neural binding by investigating neural synchrony across different frequency bands. This study examined the possibility that dysregulated alpha-gamma modulation may be crucially involved in aberrant brain functioning in autism spectrum disorder (ASD). Magnetoencephalographic data were recorded from 13 adult participants with ASD and 16 controls. The time-coursed sources averaged over a primary visual area 1 and fusiform gyrus area were reconstructed with the minimum-norm estimate method. The alpha-gamma PAC was further calculated based on these sources. The statistical analysis was implemented based on the PAC and directed asymmetry index. The results showed the hyper-activity coupling for ASD at the no-face condition and revealed the importance of alpha-gamma phase modulation in detecting a face. Our data provides novel evidence for the role of the alpha-gamma PAC and suggests that the globe connectivity may be more critical during visual perception.

## Introduction

Autism spectrum disorder (ASD) is a neurodevelopmental disorder characterized by repetitive behaviors with restricted interests and usually accompanied by mental retardation, epilepsy, hyperactivity, withdrawal, and other emotional disorders (American Psychiatric Association, 2013). According to the latest report of the US Centers for Disease Control and Prevention, 1.47% of 8-year-old children have symptoms of ASD. The diagnosis of ASD is obtained through clinical observation and testing of social disorders, language disorders, and intractable behaviors. However, these traditional psychiatric approaches are limited in some cases. Therefore, new biomarkers discovered by neuroscience approaches are critical to improving our understanding of ASD and helpful for the diagnosis of ASD.

Visual perception has been demonstrated to be abnormal in ASD with different neuroscience approaches based on brain oscillations (Stroganova et al., 2012; Sun et al.,2012; Peiker I et al., 2015). Brain (or neuronal) oscillations represented by rhythmic or repetitive patterns constitute an essential part of the brain’s electrophysiological activities. The oscillatory pattern implies brain function generated from a synchronized network linking single-neuron activity to behavior (Buzsáki and Draguhn, 2004). Recent research has discovered that gamma oscillation is abnormal in ASD during visual processing (Sun et al.,2012; Khan et al., 2013; Mamashli et al., 2018). The power of alpha oscillations decreased during visual processing and was associated with the dynamics of large-scale networks (Zhang Y et al., 2019). In our previous study, we found that gamma oscillations were deficits in adults with ASD during visual perception, suggesting a potential biomarker for the diagnosis of ASD which uses gamma oscillatory changes and the response time as a neurophysiological index and a behavioral index respectively (Sun et al.,2012). Unfortunately, we did not investigate cross-frequency coupling (CFC) to understand the underlying mechanism of interactions of different frequencies.

Phase-amplitude coupling (PAC) is one of the best study forms of CFC, in which the phase of lower-frequency oscillation modulates the amplitude of high-frequency oscillation (Seymour et al., 2017). The measure of PAC is specific to a single cortical area but also provides a direct estimation of large-scale networks and local processing interfaces. PAC became a crucial approach to understanding the cortex-cortex mechanism and is also considered a general mechanism framework for timing communication between neuronal networks (Bergmann and Born, 2018).

Recent studies have focused on the alpha-gamma phase-amplitude coupling in the context of the visual system (Voytek et al., 2010; Spaak et al., 2012; Bonnefond and Jensen, 2015). These studies have found that alpha phases have a distinct effect on gamma oscillations through local excitatory and inhibitory interactions (Buzsáki and Wang, 2012). Furthermore, the new concept of a unified framework based on nested oscillations, which interprets how the modulation of the local interaction is involved in the selective routing of information in cognitive networks, was established (Bonnefond et al., 2017). It has been reported that individuals with ASD exhibited reduced alpha-gamma PAC during visual grating tasks (Seymour et al., 2019) and facial recognition tasks (Khan et al., 2013; Mamashli et al., 2018). There is an assumption that the atypical patterns of PAC in ASD are related to dysregulation of connectivity during sensory processing (Kessler et al., 2016; An et al., 2021). Although a few studies mentioned by Seymour et al. (Seymour et al. 2017) verified the existence of atypical cortical connectivity in ASD, we still lacked knowledge about the mechanism of communication between different brain areas.

In this study, we recorded MEG data from adults with ASD and controls. We analyzed the data using alpha-gamma PAC and directed connectivity measures to investigate alpha-gamma modulation and feedback/feedforward connectivity.

## Materials and Methods

### Participants

We utilized the same dataset in our previous study. There were 13 participants with ASD (11 males and 2 females, mean age: 30.3 years; range: 20-44 years) and 16 healthy controls (12 males and 4 females, mean age: 29.7 years; range: 19 - 46). The clinical diagnosis was made for participants with ASD (details in Sun et al., 2012). Each participant should give written informed consent. The study was approved by the ethics committee of the Goethe University (Frankfurt, Germany) and conducted by the ethical standards set by the Declaration of Helsinki.

### Stimuli and task

A random sequence of Mooney stimuli, consisting of 60 upright and 30 inverted-scrambled stimuli, was presented for each run of the visual perception experiment (Fig.1A). Each stimulus lasted 200 ms. The inter-stimulus interval ranged between 3500 and 4500ms. Each participant with a hand assignment counterbalanced over all participants should respond to each stimulus with an interactive device and the responses were recorded on the presentation computer. At least four experimental runs were involved for each participant.

**Figure 1.**
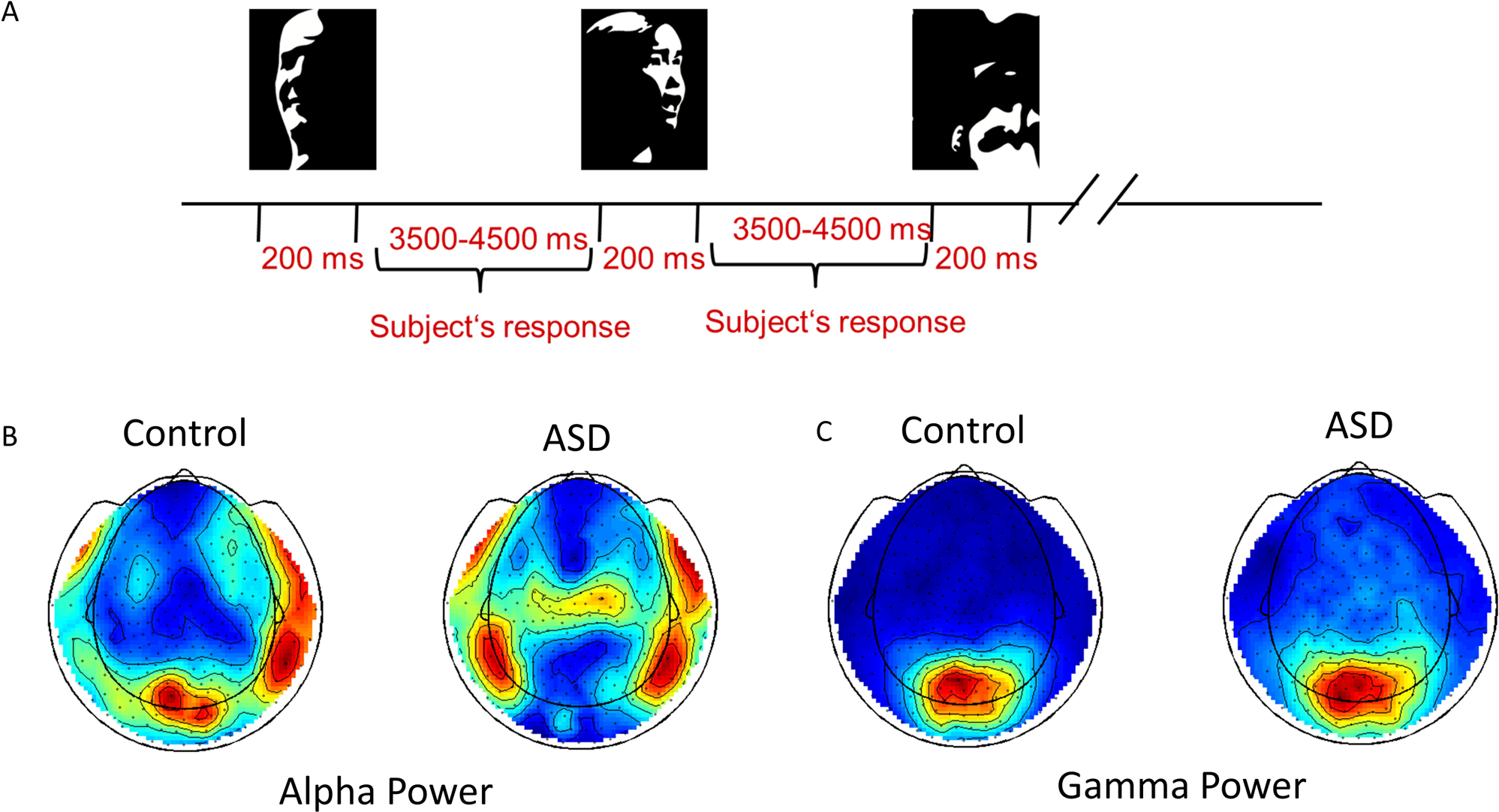
Visual Perception: (A) Paradigm and (B) interested alpha and gamma powers.

### MRI data acquisition

Structural MR images were obtained from a 3T Siemens Allegra scanner with a head coil. The protocol of the 3D MPRAGE sequence (160 slices, voxel size = 1 x 1 x 1 mm, FOV = 256, TR = 2300, TE = 3.93) was used during scanning. The positions of the nasion and 1 cm anterior to the tragus of the left and right ear were marked with vitamin-E pills to co-register the MEG and MRI data during the source localization.

### MEG data acquisition

A 275-channel whole-head MEG system (Omega 2005; VSM MedTech) was used in this study for MEG signals acquisition. A bandpass fourth-order Butterworth filter was preset with a low-frequency cutoff of 0.5 Hz and a high-frequency cutoff of 150 Hz. MEG signals were recorded with a 600 Hz sample rate. The head position was re-localized before each run to guarantee that the head movement was less than 5 mm along three directions. Runs with strong movements were discarded. Behavioral responses were recorded via a presentation computer. The differences in head coordinates between participants and groups have been examined. They were not significant and less than one electrode space of Euclidean distance for a typical EEG system.

### MEG data pre-processing

The MEG data was pre-processed using the FieldTrip Toolbox (Oostenveld et al., 2011). The trials were defined from the continuously recorded MEG from −1000 to 1000 ms concerning the onset of visual stimuli related to the face and no-face conditions. Only data with correct responses was reserved. Others were discarded. The data epochs contaminated with noises such as eye blinks, muscle activity, or jump artifacts were removed. The automatic artifact detection and rejection routines provided by the FieldTrip software were used during the trial detection. The triggers detected with the Fieldtrip routines were imported into the Brainstorm software (Tadel et al., 2011) for trial averaging. The averaged trial was filtered with a bandpass 3rd order filter with cutoff frequencies of 1Hz and 120Hz and corrected with baseline (from −500ms to −100ms). The filtered data was further used for source localization.

### Source localization

Each participant’s MRI was segmented using freesurfer software (Dale et al., 1999) and imported into Brainstorm as the head model. The source localization was performed by the minimum-norm estimate (MNE) method (Hämäläinen and Ilmoniemi, 1994) in Brainstorm. This technique provides a method to estimate the source power over the cortical surface using MEG sensors. Before initiating the analysis, noise covariance and data covariance were both estimated.

### Region of interest (ROI) definition

Two specific regions were chosen to measure the connectivity within the visual perception system. The first region, primary visual area 1 (V1), was identified using the Brodmann atlas (Brodmann, 1909). The second region, the fusiform gyrus area (FG), was selected using the Desikan-Killiany atlas (Desikan et al., 2006). The areas of V1 and FG were anatomically separated, as shown in Figure 2. Time-coursed sources from these regions of interest were extracted and averaged over each region individually.

**Figure 2.**
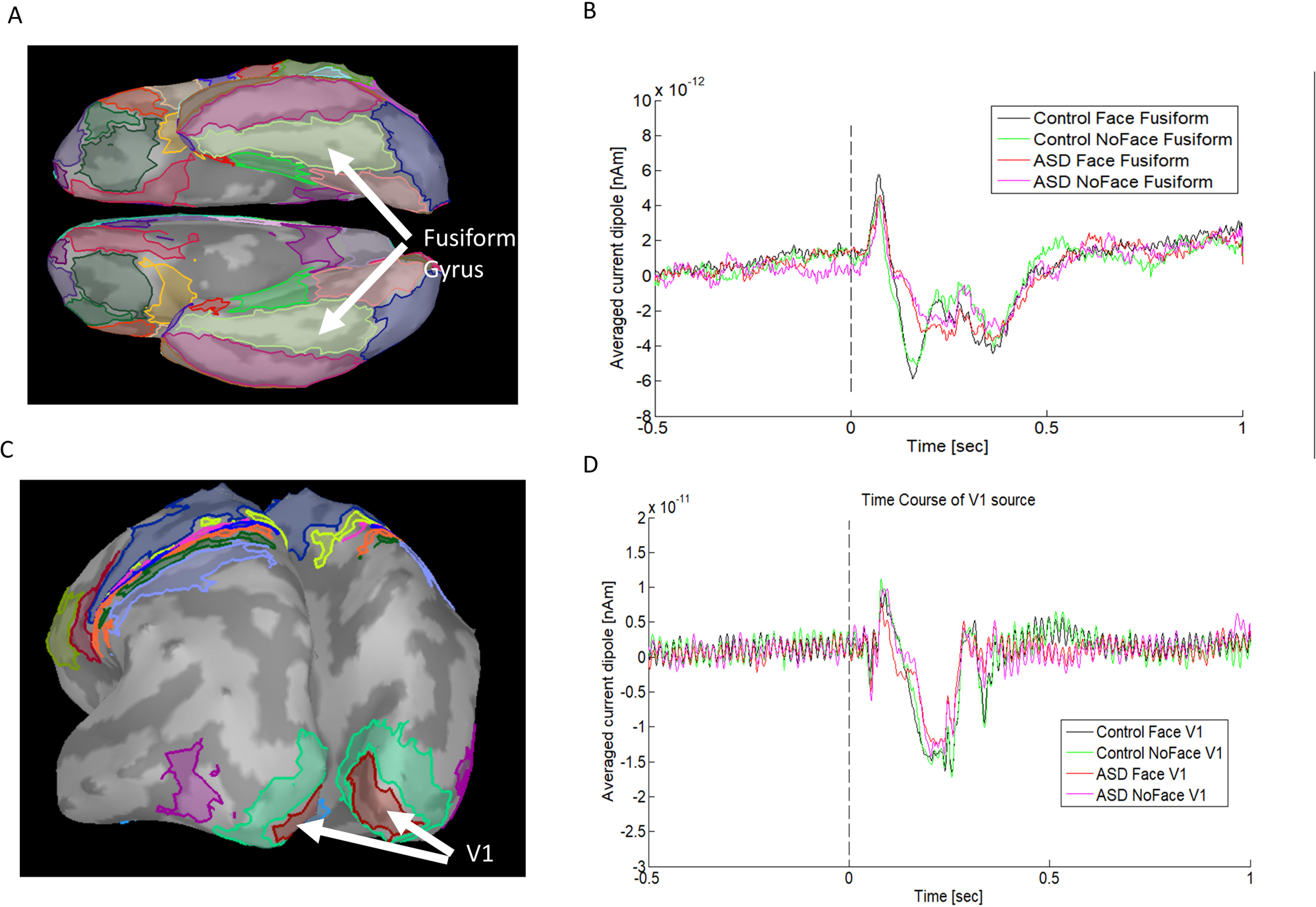
The time-coursed sources averaged over ROIs from one subject. (A) Fusiform gyrus area; (B) the time-coursed sources averaged over Fusiform Gyrus areas on the left and the right hemispheres; (C) primary visual area 1 (V1); (D) the time-coursed sources averaged over the V1s on the left and the right hemisphere.

### Phase amplitude coupling

The study examined the sources of V1 and FG over time to detect any changes in alpha-gamma phase-amplitude coupling (PAC). Brainstorm was used to compute the PAC values for phase frequencies ranging from 7-13Hz and amplitude frequencies from 34-120Hz, starting from stimulus presentation (0.0ms) to 1000ms. The phase step size was 1Hz. The amplitude-frequency step size was 2Hz. To determine which gamma frequencies had the most significant changes in PAC, Seymour et al. (2017) used a range of amplitude frequencies (34-120 Hz). We selected the same frequency of 34Hz as the lower limit of the range to meet the reasonable detectable amplitude frequency requirement for alpha phases.

### Statistical analysis for PAC

This study used PAC values computed from the time-coursed sources of V1 and FG to do the statistical analysis. A non-parametric 2×2 ANOVA with permutations was performed for each region with two factors: group (controls vs. ASD patients) and condition (face vs. no-face). This analysis aimed to investigate the main effects and interactions of the factors. The statistical results were corrected for the multiple-comparison problem in phase frequency and amplitude frequency using the Holm statistical analysis with 5000 permutations. An alpha level of 0.05 was set.

### Directed connectivity for V1 and FG

To measure directed functional connectivity, we employed a bivariate Granger causality method (Granger, 1969; Dhamala et al., 2008) based on the time-coursed V1 and FG sources (0–1.0 s after stimulus onset). The values were averaged for V1 and FG individually and then calculated for each pair of interests. These values were then used to indicate the strength of connectivity from one region (V1 or FG) to the other (FG or V1).

The Directed Asymmetry Index (DAI) has been used to measure the asymmetries of directed connectivity (Bastos et al., 2015b). This DAI (equation. 1) is used to determine whether the Granger causality influence is feedforward (DAI>0) or feedback (DAI<0), as noted by Seymour et al. (2019). The DAI values were compared statistically between different groups.

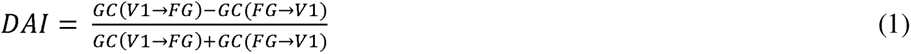

Where *GC*(*V1→FG*) means the granger causality value is calculated from V1 to FG and *GC*(*V1→FG*) means the granger causality value is calculated from FG to V1.

### Statistical analysis for connectivity

The regions of interest are the left V1 (V1 L), the right V1 (V1 R), the left FG (FG L), and the right FG (FG R). A connectivity map based on Granger causality among these four regions was analyzed statistically using the non-parametric 2×2 ANOVA method similar to the one mentioned above.

### Behavioral correlations with PAC and connectivity

To investigate the relationship between alpha-gamma PAC and behavior, we computed correlations between each alpha-gamma PAC, reaction times, detection rates, and the discrimination index (A’) for both the ASD and control groups for the face and no-face conditions separately. We obtained correlations for each alpha-gamma PAC at different conditions with reaction times, detection rates, and the discrimination index A’. We also computed correlations between connectivity and behavior data. Significant correlations were found among regions (V1 L, V1 R, FG L, and FG R) at different conditions with reaction times, detection rates, and the discrimination index A’. Only significant correlations that met an alpha level of 0.05 with Bonferroni correction (Curtin and Schulz, 1998) were considered.

## Results

### Alpha band and Gamma band

The oscillatory powers of alpha (7-13Hz) and gamma (34-120Hz) were calculated at the sensor level. For both control and ASD groups, this study found that the strong gamma power was located at the occipital lobes (as shown in Fig. 1B) and the strong alpha power was located at the parietal lobes. Interestingly, the control group also had strong alpha power at the occipital lobes.

### Sources of ROIs

Two regions of interest (ROIs) were chosen in each hemisphere. The time-coursed sources were reconstructed using the MNE approach, as shown in Figure 2B and Figure 2D. These changes revealed consistent patterns of hierarchical connectivity. The connectivity between V1 and FG (V1 - FG) was feedforward, while the connectivity between FG and V1 (FG - V1) was feedback (Bastos et al., 2015a,b; Michalareas et al., 2016).

### ANOVA analysis for Alpha-gamma PAC

This study examined changes in alpha-gamma PAC based on the source activities from the V1 and FG areas of the brain. The ANOVA analysis within V1 showed that the main effects of the group were statistically decreased at 11-12Hz phase frequencies and 30-40Hz amplitude frequencies, indicating that the alpha modulation in the ASD group was strong (Fig. 3A). The analysis also revealed a main effect of condition at 10-12Hz phase frequencies and 48-60Hz amplitude frequencies, suggesting that the phase-amplitude modulation was downregulated for responses to face vs no-face stimuli (Fig. 3B). Furthermore, the interaction between the factors group × condition showed that PAC activity in controls at 8-10Hz phase frequencies and 30-32Hz amplitude frequencies was significantly upregulated in responses to face-stimuli relative to ASD-patients (Fig. 3C). It’s noteworthy that the main effect of condition was found in FG only at the higher amplitude frequency with the centralized amplitude frequency of 100 Hz modulated by 11-12Hz phase frequencies (Fig. 3D).

**Figure 3.**
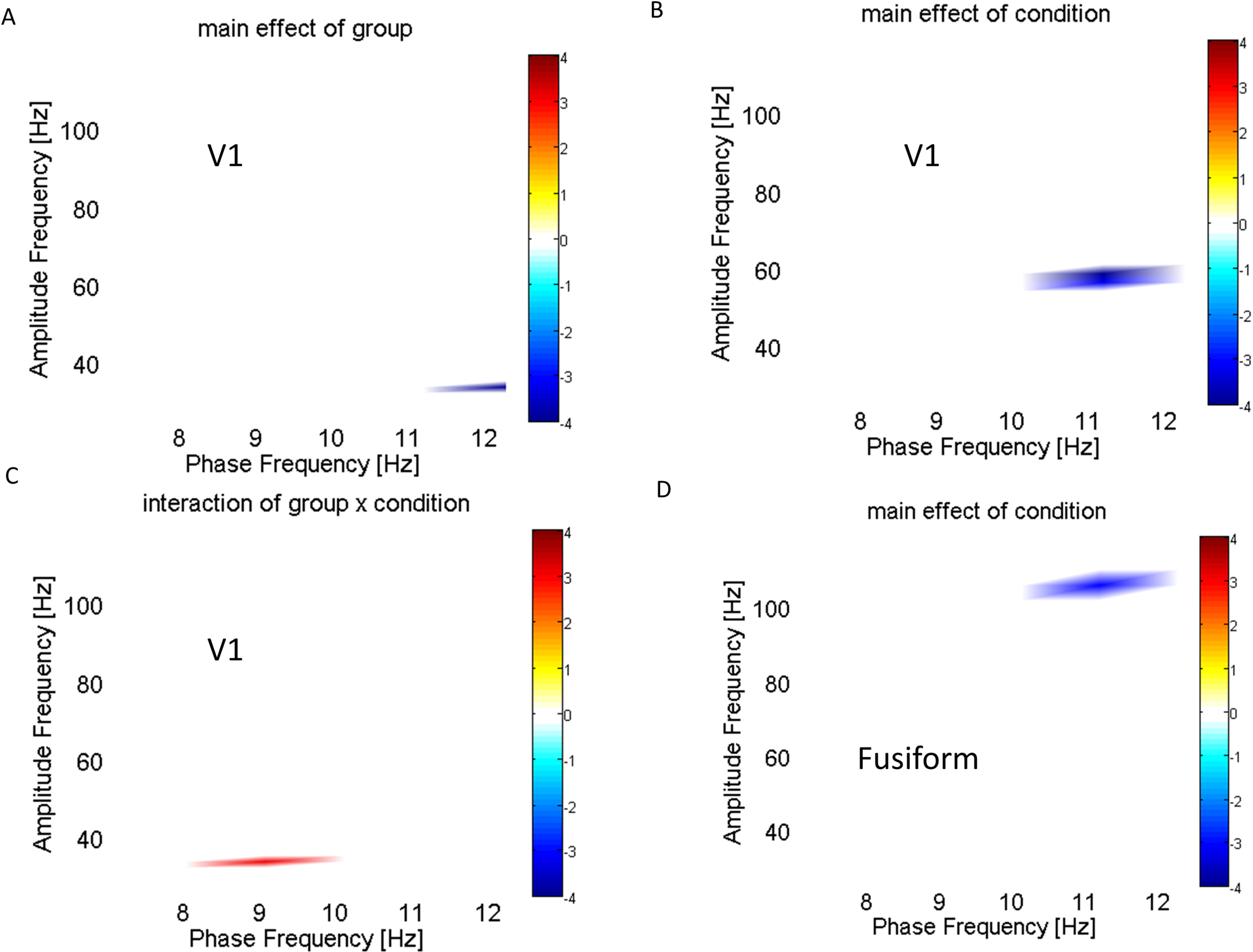
ANOVA analysis for alpha-gamma PAC: (A) Main effect of Group in V1; (B) Main effect of condition in V1; (C) the interaction between the two factors (control vs. ASD and face vs. no-face); and (D) Main effect of condition in FG. For the main effect of groups in (A) and (D), the blue colors indicate the increased PAC in the ASD group. For the main effect of condition in (B) the blue color indicates the higher alpha-gamma PAC to no-face stimuli. In C, the red color indicates higher alpha-gamma PAC in the face condition for controls vs. autistic patients. The statistical analysis is multiple-comparison corrected and the alpha level is set to 0.05.

### Granger Causality for connectivity

We used Granger causality to measure the directed functional connectivity between different parts of the brain (V1 L, V1 R, FG L, and FG R). We found that the connectivity was increased in patients with ASD with face condition for V1 L-to-FG L, V1 R-to-FG R, FG L-to-V1 L, and FG R-to-V1 R (Fig. 4). The prominent increased connectivity was presented from V1 to FG as well as from FG to V1 in the right hemisphere. There were increased connectivities between V1 and FG in the local network and between the right and left fusiform gyrus in the global network. The statistical analysis demonstrated significant connectivity of the main effect of the group for FG R-to-FG L, indicating a synchronized oscillation between the right and left hemispheres (see Figure 5A). The main effect of the condition revealed the significant connectivity for V1 L-to-FG L and FG L-to-FG R, indicating that the Granger causality is more robust in response to face stimuli compared to no-face stimuli (Fig. 5B). Moreover, the interaction between the factors (group and condition) showed that in controls, the connectivity for FG L-to-V1 R was increased significantly in response to face-stimuli compared to ASD-patients (Fig. 5C). We employed the DAI metric to measure asymmetries in feedforward and feedback connectivity for both groups and conditions. In the left hemisphere, the control group showed a greater flow in feedforward connectivity (positive DAI values) at the no-face condition. Conversely, in the right hemisphere, the control group showed an increased flow in feedback connectivity (negative DAI values) at the face condition (Fig. 6).

**Figure 4.**
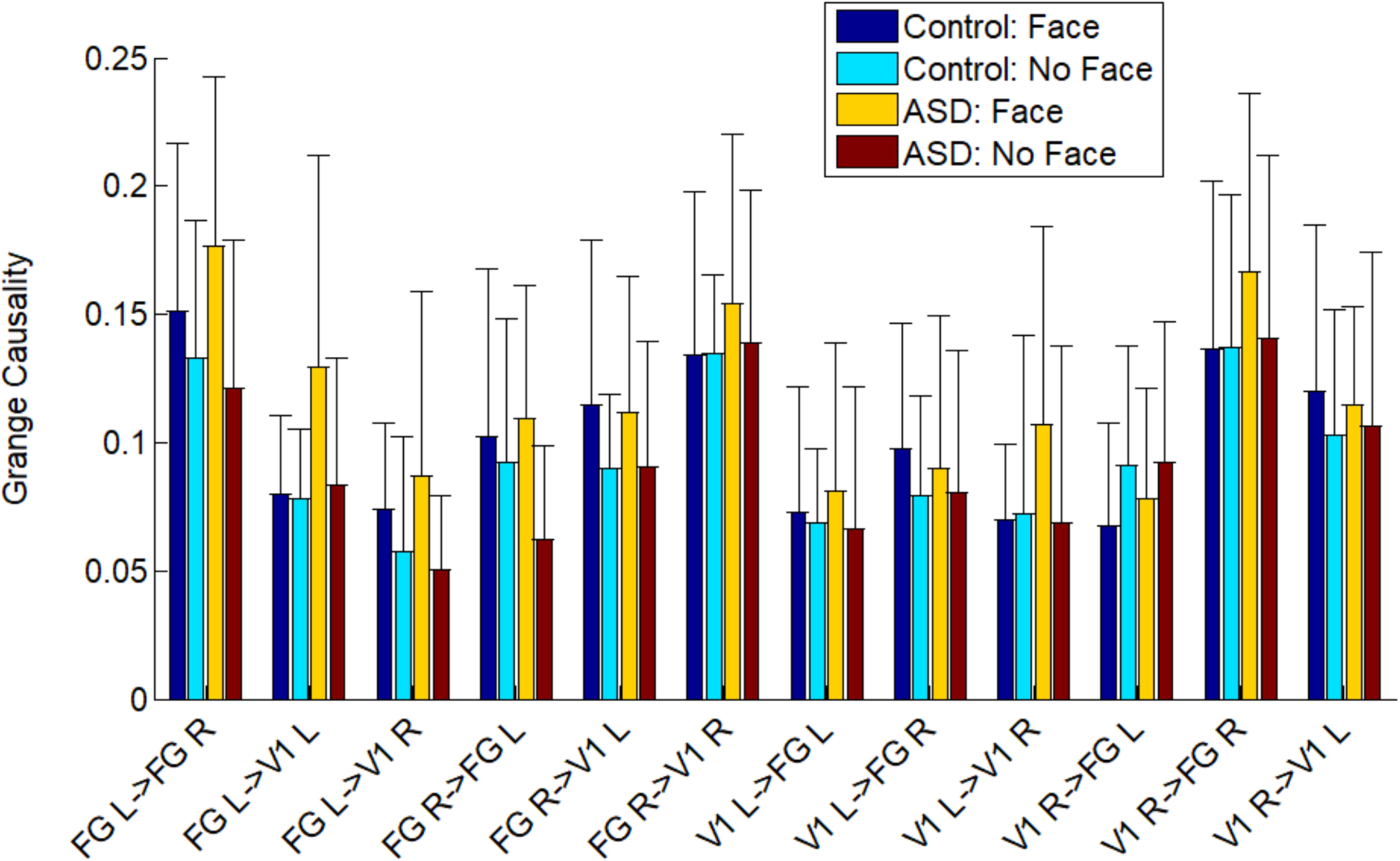
Grange causality values for V1-FG directed connectivity were computed based on the groups and the conditions. Both the local connectivity and the global connectivity were estimated.

**Figure 5.**
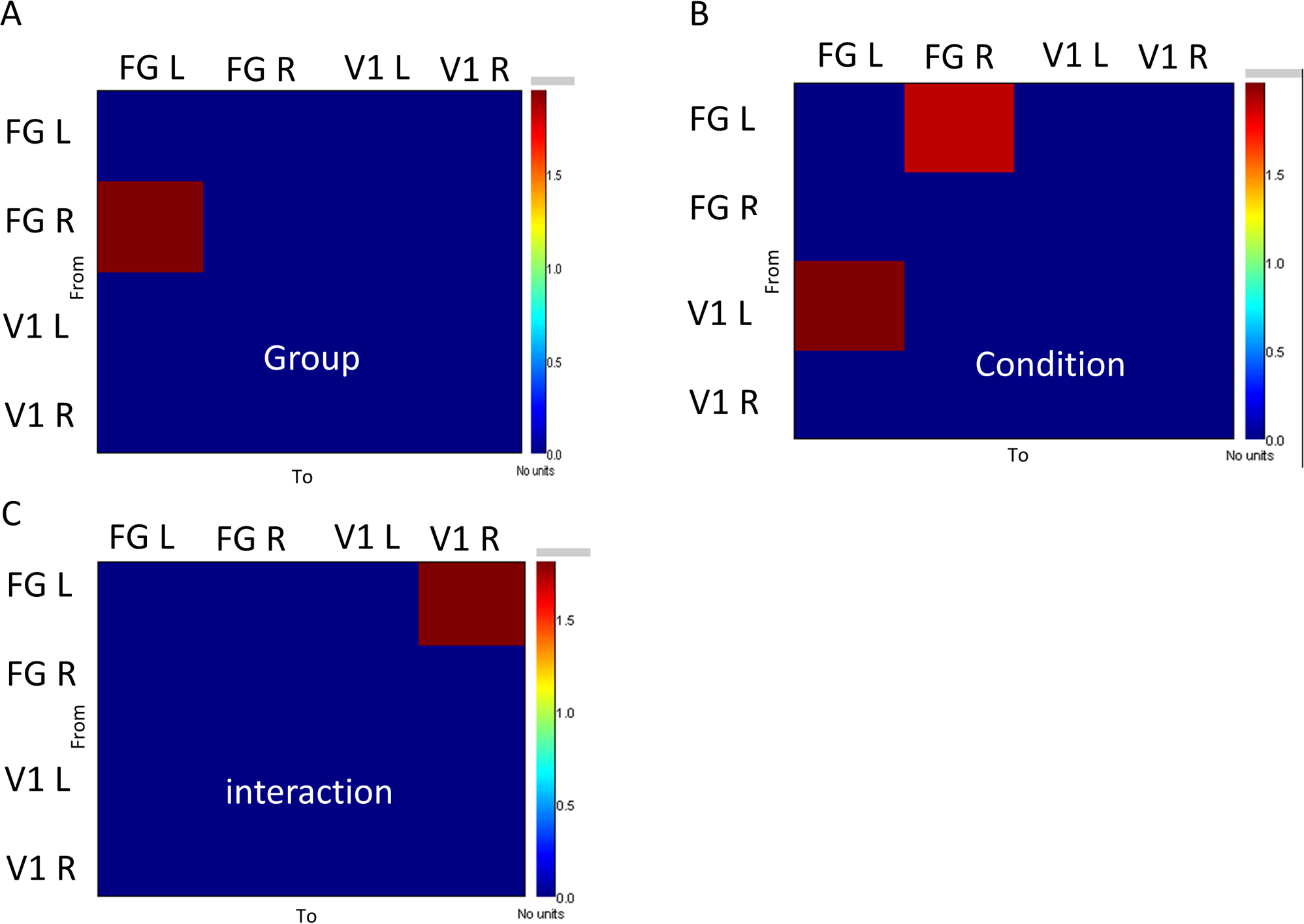
ANOVA analysis was implemented based on the grange causality values in Figure 4: (A) the Main effect of the group, (B) the Main effect of the condition, and (C) the interaction between the two factors (control vs. ASD and face vs. no-face). For the main effect of the group, the red color indicates the stronger connectivity in controls. For the main effect of the condition, the red colors indicate the stronger connectivity to face stimuli. For the interaction between the two factors, the red color suggests higher connectivity in the face condition for control vs. autistic patients. The statistical analysis is multiple-comparison corrected and the alpha level is set to 0.05.

**Figure 6.**
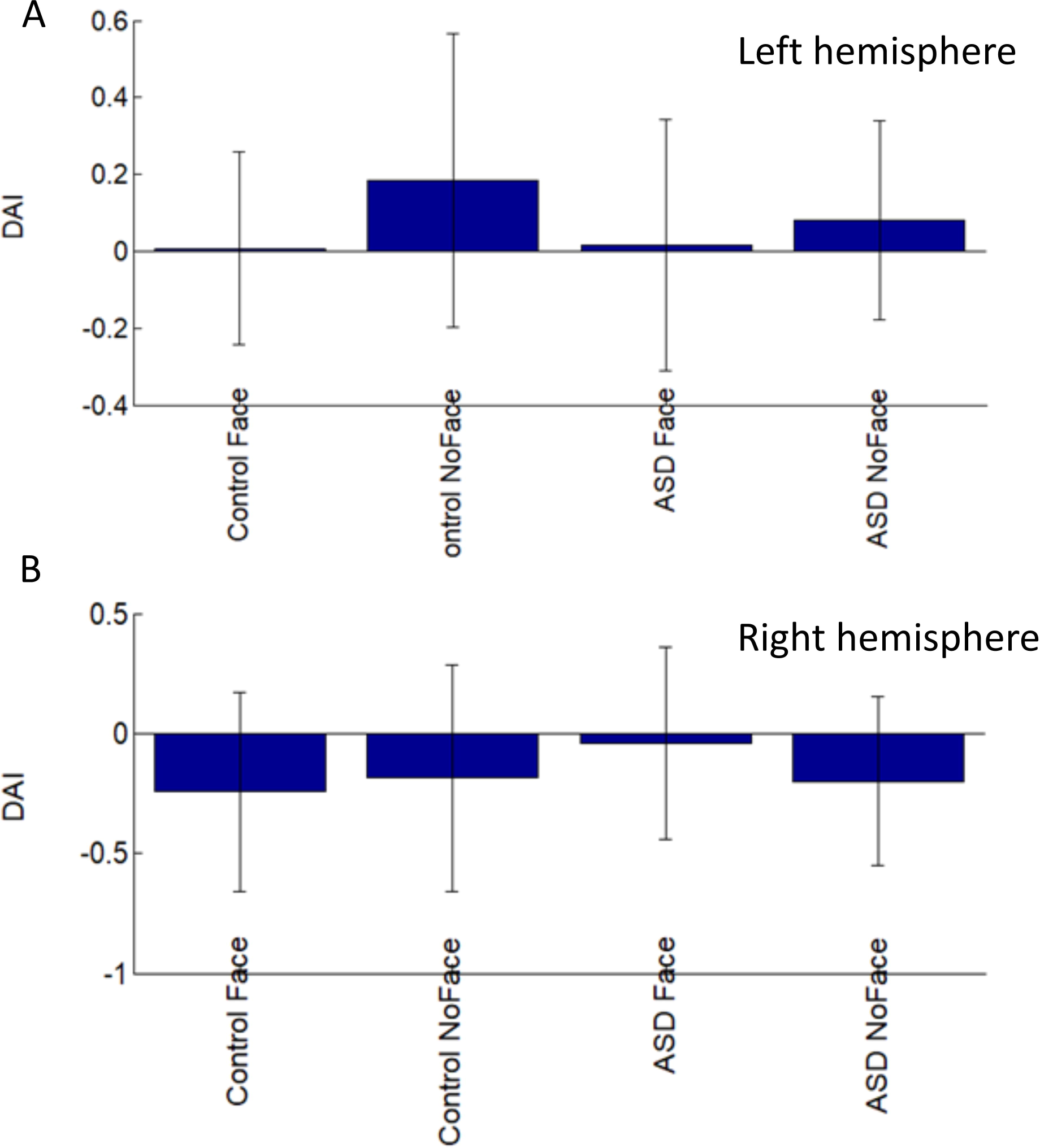
The directed asymmetry index (DAI) was computed for each group and each condition individually at the left hemisphere (A) and the right hemisphere (B).

### Behavior correlation analysis

Table 1 and Table 2 showed the correlation between behavioral data for control and ASD groups with alpha-gamma PAC. There was a significant positive correlation between the PAC in V1 and the detection rate of the control group at the face condition. Additionally, there were significant negative correlations between the PAC in V1 and the detection rate for the control group at the no-face condition and the ASD group at the face condition. There was a positive correlation between PAC in FG and the detection rate in the control and ASD groups at the no-face condition. However, there was a negative correlation between PAC in FG and the detection rate in the ASD group at the face condition. In the control group, no significant correlation was found for the index A’ at the face condition, while a significant positive correlation was observed at the no-face condition. However, the ASD group showed a significant negative correlation for the index A’ at both conditions in V1. Only one significant negative correlation was observed between the index A’ and the PAC in FG at the face condition for the control group. For the reaction time, the control group displayed a significant negative correlation in V1 at both conditions, while the ASD group showed a significant positive correlation only at the face condition. In FG, the reaction time showed a negative correlation at the face condition but a positive correlation at the no-face condition for the control group. For the ASD group, the reaction time showed a positive correlation with the no-face condition.

**Table 1.**
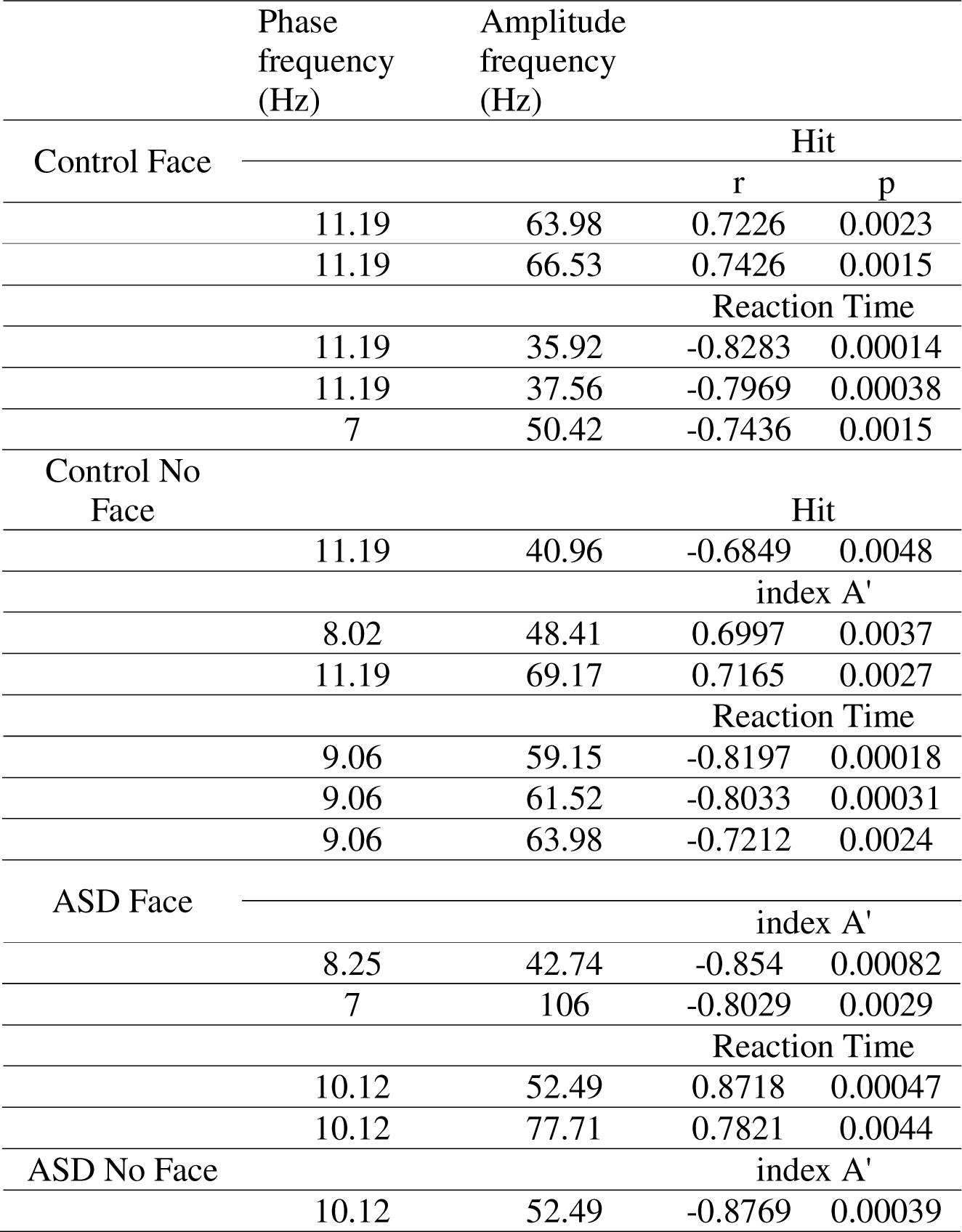
Correlation between behavior and PAC in V1.

**Table 2.**
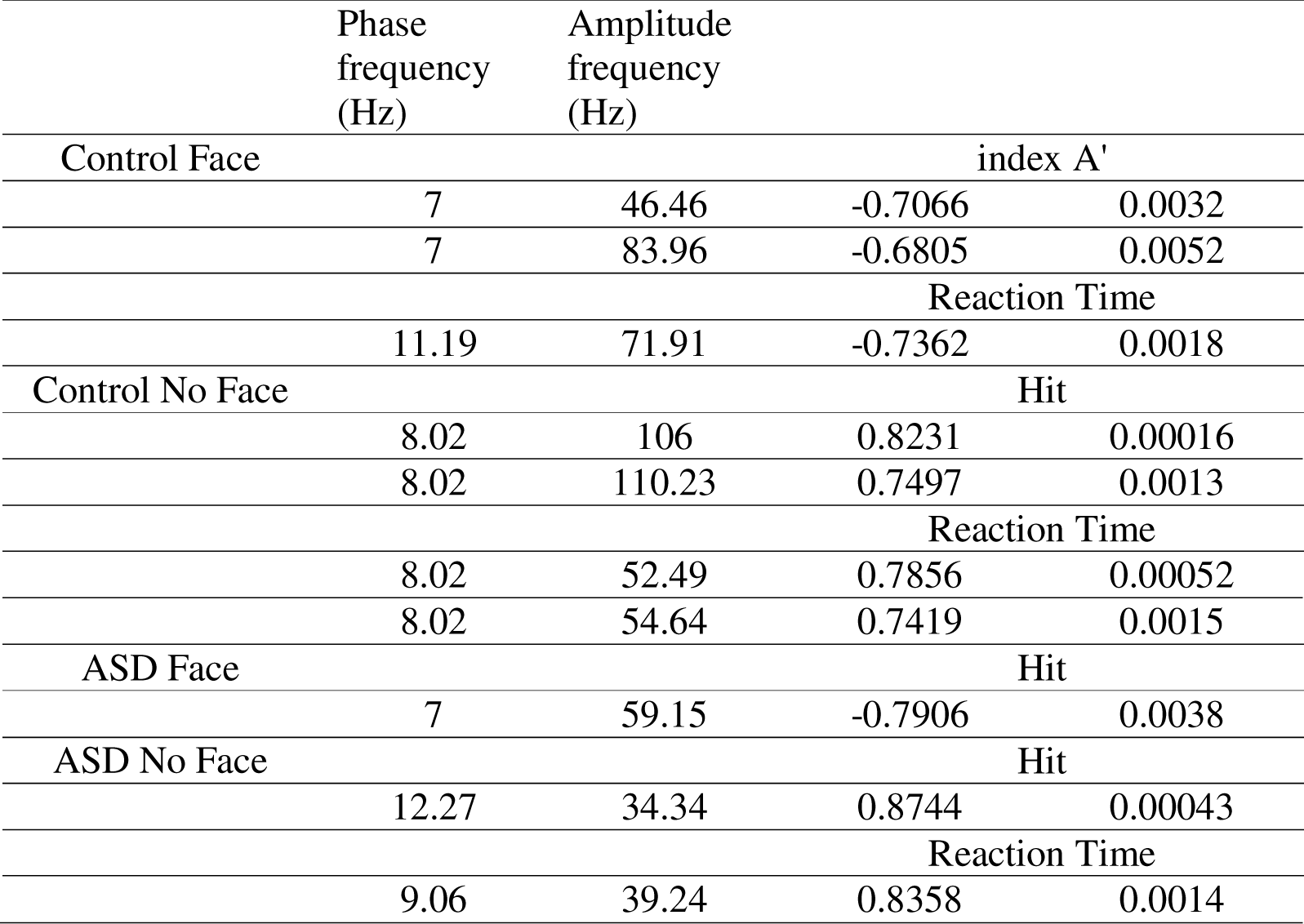
Correlation between behavior and PAC in FG.

|  | Phase<br>frequency<br>(Hz) | Amplitude<br>frequency<br>(Hz) |  |  |
| --- | --- | --- | --- | --- |
| Control Face |  |  | index A' |  |
|  | 7 | 46.46 | -0.7066 | 0.0032 |
|  | 7 | 83.96 | -0.6805 | 0.0052 |
|  |  |  | Reaction Time |  |
|  | 11.19 | 71.91 | -0.7362 | 0.0018 |
| Control No Face |  |  | Hit |  |
|  | 8.02 | 106 | 0.8231 | 0.00016 |
|  | 8.02 | 110.23 | 0.7497 | 0.0013 |
|  |  |  | Reaction Time |  |
|  | 8.02 | 52.49 | 0.7856 | 0.00052 |
|  | 8.02 | 54.64 | 0.7419 | 0.0015 |
| ASD Face |  |  | Hit |  |
|  | 7 | 59.15 | -0.7906 | 0.0038 |
| ASD No Face |  |  | Hit |  |
|  | 12.27 | 34.34 | 0.8744 | 0.00043 |
|  |  |  | Reaction Time |  |
|  | 9.06 | 39.24 | 0.8358 | 0.0014 |

Behavioral data from both control and ASD groups were analyzed for correlation with connectivity. We found a significant negative correlation (Fig. 7A, r = −0.586, p = 0.0215) between detection rate and DAI in the right hemisphere for the control group at the face condition, indicating that feedback connectivity (negative DAI values) increases with higher detection rate. In addition, a significant positive correlation (Fig. 7B, r = 0.609, p=0.0468) between detection rate and DAI in the left hemisphere of the ASD group was also found, suggesting that the autistic patients had a higher detection rate but decreased feedback connectivity. However, there were no significant correlations between behavior and the index A’ and the reaction time.

**Figure 7.**
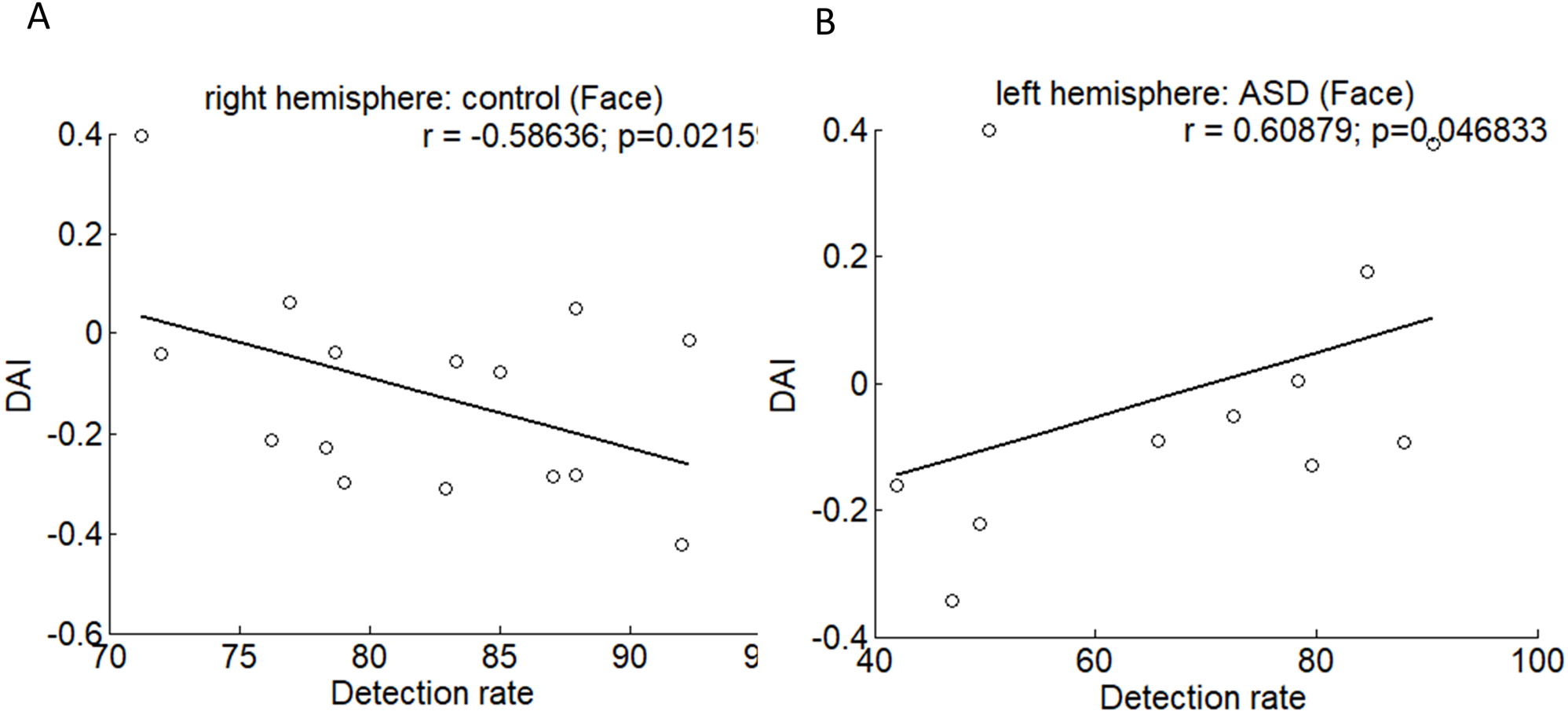
Correlations between behavior and DAI. For the control group, the correlation between DAI and the detection rate was negative at the face condition (A). For the ASD group, the correlation between DAI and the detection rate was positive at the face condition (B).

## Discussion

We investigated the oscillation-based functional connectivity by visual perception experiments with autistic patients and age-matched control subjects. Our research has shown that there is an increase in Granger causality influence from FG L to V1 L and FG R to V1 R in individuals with ASD (Fig. 4). Furthermore, we have observed a decrease in both feedback and feedforward connectivity in the no-face condition in the ASD group as compared to the control group (Fig. 6). Additionally, there is an increase in alpha-gamma coupling in V1 (Fig. 3A), measured by PAC, indicating dysregulation of connectivity in the visual perception system of adult patients with autism. Our findings contradicted previous findings of decreased connectivity and coupling in autistic adolescents for visual gating stimuli (Seymour et al., 2019).

### Phase amplitude coupling

Our study showed a significant reduction in alpha-gamma PAC in the primary visual area V1 for the group effect (Control vs. ASD) and the condition effect (face vs. no-face) (refer to Fig. 3A and 3B). The statistical results obtained from ANOVA revealed hyper-activity coupling at the no-face condition for ASD, consistent with some M/EEG studies (Orekhova et al., 2007; Cornew et al., 2012). Interestingly, the interactive analysis showed that the relative changes between the face and no-face conditions revealed a deficit in modulation for ASD.

In recent studies, PAC has been used as a reliable indicator for the imbalance between excitatory and inhibitory neurons in ASD. While other studies found a reduction in PAC in ASD, our study found an increase in PAC in ASD cases. Therefore, we assumed that the PAC beyond an acceptable range could lead to an imbalance between excitatory and inhibitory neurons. Abnormal PAC can result from low or high-frequency ‘noisy’ activity in local inhibitory processes, disrupting visual perception.

PAC is assumed to be a general cortical mechanism for oscillatory multiplexing to link connectivity at the global and local scales (Canolty and Knight, 2010; Seymour et al., 2017, 2019). Researchers have found a strong correlation between PAC and subjects’ performance in tasks such as detection rate, index A’, and reaction time. In controls, the strong ability of alpha phase modulation resulted in improved reaction time. However, PAC was negatively correlated with the ability to discriminate in V1 in individuals with ASD, indicating dysregulated visual perception functions.

### Feedforward/feedback connectivity

We focused on the connectivity between two specific brain areas, V1 and FG, involved in face recognition and processing. We estimated the connectivity for V1-to-FG in each hemisphere instead of taking the whole visual area. The asymmetries in V1-FG connectivity were observed at the source level. In ASD, we found feedback connectivity (FG L -> V1 L) stronger than feedforward connectivity (V1 L->FG L) in the left hemisphere but not in the right hemisphere. This finding aligns with our PAC results. The globe feedback/feedforward connectivity in this study was examined using measures of directed functional connectivity (e.g., Granger causality) in concert with higher-level cognitive tasks that involve a more extended set of cortical regions.

During the visual perception of the mooney face recognition, we observed a slight increase in connectivity from V1 L-to-FG L and V1 R-to-FG R at both face and no-face conditions in ASD (Fig. 4), suggesting un-equivalent levels of feedforward information flow during the visual perception processing between groups. This finding indicated that the autistic participants were overmodulated over the interneurons. This unbalanced information flow implied that the impaired feedback and feedforward information flow strongly affected the discrimination of face recognition.

Although there was no significant difference between groups and conditions for DAI for both hemispheres, a phenomenon arose, indicating that the mean DAI at the no-face condition was larger in control, which suggested a reduction in the feedforward flow of information from lower to higher visual regions in ASD (Fig. 6A). We found a similar reduction for the feedback flow of information from higher to lower visual regions in the right hemisphere in ASD (Fig.6B). Therefore, asymmetries occur not only in flow directions but also in different hemispheres. This finding suggests that the reduced modulation ability in both directions of visual information caused atypical visual processes in ASD.

We conducted a global connectivity check for V1 and FG in each hemisphere. Based on ANOVA analysis (control vs. ASD; face vs. no-face), we did not find any significant connectivity in the local network. However, we found strong connectivity significantly in the global network. This evidence suggested that global oscillation synchronization might be crucial in visual perception processing.

Our study revealed significant correlations between DAI and the detection rate for the face condition in both the control and ASD groups. We found significant correlations in the right hemisphere in the control group and the left hemisphere in the ASD group. It further implicated that unbalanced feedback connectivity was critical for face detection.

### Limitations

Firstly, the behavior performance assessments only focused on detection rate, index A’, and reaction time. We realized that it would be better to have more other assessments, such as GSQ scores. Secondly, there was a limited number of participants in each group in our study. Thirdly, only a few visual areas were selected for the connectivity investigation.

## Conclusions

Our data provided novel evidence for the roles of alpha-gamma PAC and connectivity in visuo-perceptual dysfunctions in ASD by demonstrating the hyper-activity PAC in ASD for the no-face condition and the reduced feedback connectivity in ASD for the face condition in the right hemisphere. This study also implied that the unbalanced global network played a crucial role in visual perception. Our data suggested that the alpha phase modulating gamma power is critical to improving behavior performance in ASD.

## Acknowledgments

The author thanks Christine Gruetzner, who collected MEG data at the Max Planck Institute for Brain Research, Frankfurt am Main, Germany, and Dr. Peter Uhlhass, who was the supervisor in the ASD project and is now a professor at Glasgow University in the UK.

## Statements and Declarations

### Data availability

The raw data are not publicly available due to ethical restrictions but the preprocessed data can be provided on reasonable request from the corresponding author.

